# Human naïve stem cell models reveal the role of FGF in hypoblast specification in the human embryo

**DOI:** 10.1101/2023.11.30.569161

**Authors:** Anish Dattani, Elena Corujo-Simon, Arthur Radley, Tiam Heydari, Yasaman Taheriabkenar, Francesca Carlisle, Simeng Lin, Jonathan Mill, Peter Zandstra, Jennifer Nichols, Ge Guo

## Abstract

The hypoblast is an essential extra-embryonic tissue set aside within the inner cell mass early in mammalian embryo development, in the blastocyst. Research with human embryos is challenging. Thus, stem cell models that reproduce hypoblast differentiation provide valuable alternatives. We show here that human naïve PSC to hypoblast differentiation proceeds via reversion to a transitional ICM-like state, from which hypoblast emerges in concordance with the trajectory in human blastocysts. We identified a window when fibroblast growth factor (FGF) signalling is critical for hypoblast specification. Revisiting FGF signalling in human embryos revealed that inhibition in the early blastocyst suppresses hypoblast formation. *In vitro*, the induction of hypoblast is synergistically enhanced by limiting trophectoderm and epiblast fates. This finding revises previous reports and establishes a conservation in lineage specification between mouse and human. Overall, this study demonstrates the utility of human naïve PSC-based models in elucidating mechanistic features of early human embryogenesis.

## INTRODUCTION

At the onset of early human embryogenesis, a fertilised zygote undergoes continuous cell division, cellular diversification, and spatial organisation to form a hollow spherical structure, the blastocyst. The blastocyst consists of three primary lineages, the epiblast, trophectoderm, and hypoblast, which will develop into the embryo proper, placenta, and yolk sac. Studies in mice have established a model of sequential lineage segregation during pre-implantation embryogenesis. This begins with the first lineage segregation, which sets apart the extraembryonic trophectoderm from the inner cell mass (ICM)^1–3^ and is followed by the second lineage segregation when the ICM differentiates into naïve epiblast and hypoblast. Morphological observation, immunostaining, and recent advances in single-cell transcriptome analysis of human embryos suggest that the sequential lineage segregation model broadly stands in human^4–6^. The segregation of trophectoderm in human begins around day 4 after fertilisation (dpf) in the morula, followed by hypoblast lineage segregation around dpf 5/ 6 in the inner cell mass of blastocyst. Cross-species comparative studies have also revealed both conserved and human-specific features of lineage specification^7–9^. However, due to limited access to human embryos, applicable approaches, and ethical considerations, knowledge of the regulatory mechanisms controlling human hypoblast lineage segregation is fragmentary.

Human pluripotent stem cells and embryo models can be valuable tools to study cell fate specification. Human naïve pluripotent stem cells (naïve PSCs) display similarity to the epiblast in 6/7 dpf blastocysts^10–13^. We found that human naïve PSCs maintained in PXGL medium retain a greater degree of developmental plasticity compared to mouse ESC counterparts^14,15^. Inhibition of MEK/ERK (PD0325901) and Activin/Nodal signalling (A83-01) enables naïve PSCs to differentiate to trophectoderm^14^. Interestingly, one day exposure to PD0325901 and A83-01 yielded a mixed cell culture comprising all three lineages in the blastocyst, including a small population of hypoblast-like cells^14^. This cellular plasticity has been exploited to form three-dimensional organoids resembling blastocysts, called blastoids, that share morphological and transcriptomic similarity to the pre-implantation embryo^16–18^. Signalling pathways involved in trophectoderm specification have been described^14,15,19^ but the naïve PSC to hypoblast differentiation path has not been properly characterised. Furthermore, a key question remains as to how the *in vitro* differentiation process relates to hypoblast lineage development in the embryo. Here, we delineate the *in vitro* naïve PSC to hypoblast differentiation trajectory and compare it with embryo lineage segregation. We also explore the role of FGF signalling, which is critical for hypoblast specification in mouse, but has previously been suggested to be less important in human.

## RESULTS

### Initial MEK/ERK and Activin/Nodal inhibition enables hypoblast differentiation from naïve PSCs

We have previously shown that human nPSCs differentiate mainly (>60%) to trophectoderm over three days continuous culture in N2B27 basal medium containing MEK/ERK inhibitor (<u>PD</u>0325901) and Activin/Nodal inhibitor (<u>A83</u>-01)^14^. In contrast, one day in <u>PDA83</u> followed by N2B27 alone yielded a mixed culture containing a small population of hypoblast cells^14^. We used this adherent nPSC differentiation to investigate the signalling requirements for hypoblast differentiation. To this end, we generated an OCT4-eGFP and SOX17-tdTomato knock-in double reporter naïve iPSC cell line, niPSC-OS1 (Figure S1A,B). niPSC-OS1 cells expressed the pluripotency marker OCT4-GFP in PXGL^10^, but few, if any, SOX17-tdTomato positive cells were observed during self-renewal (Figure S1C). SOX17-tdTomato positive cells became readily visible under fluorescence microscopy in differentiation (Figure 1A, B). We quantified reporter expression by flow cytometry over the course of the three days of differentiation in either continuous PDA83 (PA condition) or one day in PDA83 with release into N2B27 (PA-N condition) (Figure 1A). The PA condition predominantly gave rise to reporter negative cells with a minor population of OCT4 positive cells. Little or no SOX17-tdTomato expression was recorded (Figure 1C, Figure S1D). The double negative cells expressed the trophectoderm marker GATA3 (Figure S1E). In the PA-N condition, SOX17-tdTomato expression emerged after 24 hours in N2B27 (Figure S1F). Typically, after 3 days of culture in PA-N, we observed a mixed culture with ∼20% SOX17-tdTomato+, ∼60% OCT4-GFP+ and ∼20% double-negative cells (Figure 1C). Immunofluorescence (IF) and reporter expression confirmed the presence of pluripotency, hypoblast, and trophectoderm lineage markers in the three subpopulations in PA-N cultures (Figure S1G). Importantly, when plated directly into N2B27 without PDA83 exposure, the culture largely remained OCT4-eGFP positive with only ∼1% of SOX17-tdTomato+ cells (Figure 1D), indicating that exposure to PDA83 is important for both trophectoderm and hypoblast specification.

**Figure 1:**
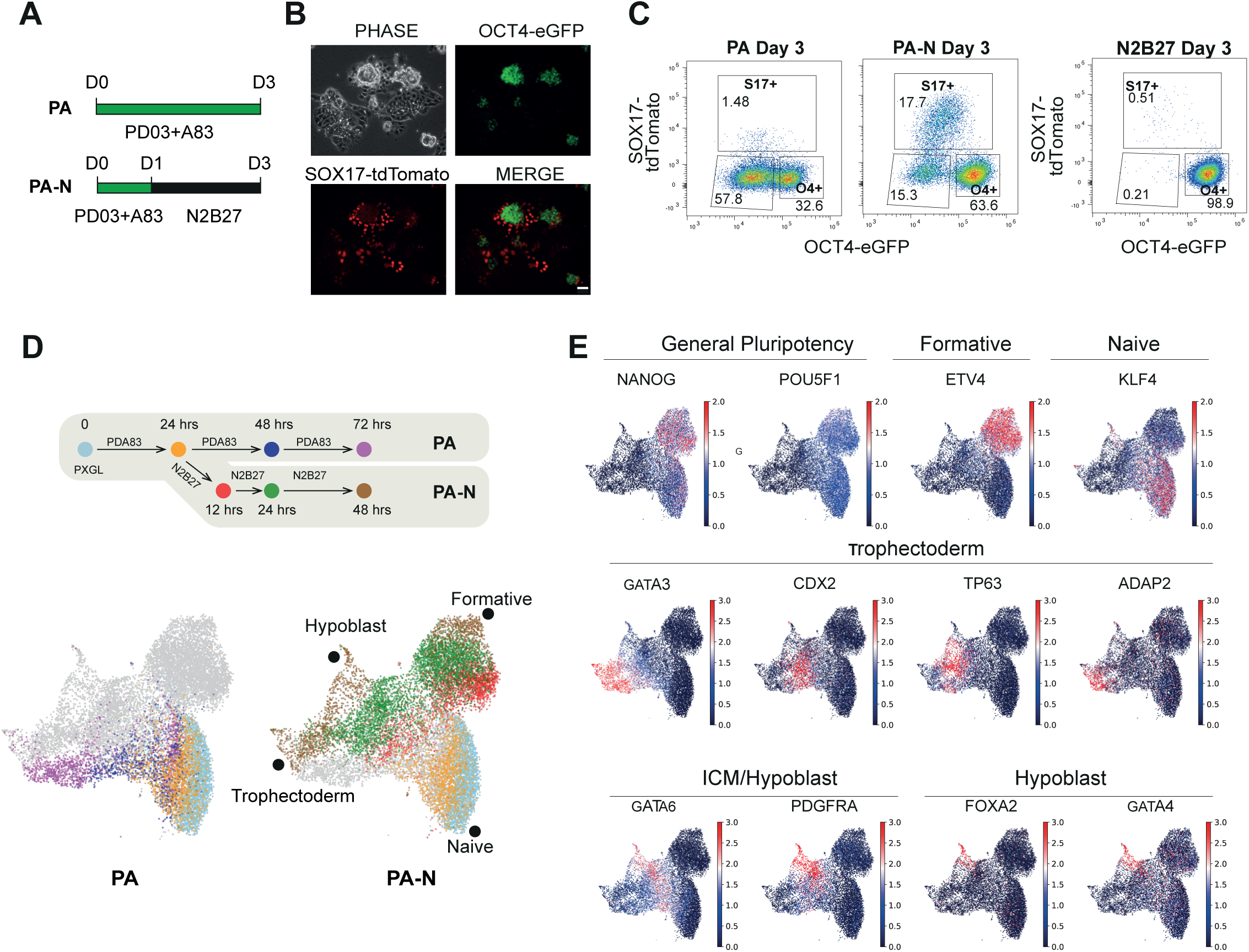
Hypoblast differentiation in human naïve iPSC-OS1 cells. (A) Schematic of nPSC differentiation in condition PA or PA-N. (B) Live image of niPSC-OS1 culture differentiated in PA-N for 3 days. Scale Bar = 50 μM. (C) Flow cytometry analysis of OCT4-eGFP and SOX17-td tomato expression after three days culture in differentiation conditions PA or PA-N. (D) Flow cytometry analysis of OCT4-eGFP and SOX17-td tomato expression after three days culture in N2B27 without prior PD03+A83 treatment. (E) Combined UMAP of the 10x single-cell sequencing of PA and PA-N differentiation time course, coloured according to condition and time. (F) Log2 expression of selected markers genes indicating differentiation trajectories in (D).

We performed single-cell transcriptome analysis across the three-day differentiation timecourse in PA and PA-N conditions to obtain information on the differentiation trajectories. Uniform manifold approximation and projection (UMAP) visualisation showed continuous progression of three distinct fates expressing trophectoderm, hypoblast or formative epiblast markers respectively (Figure 1E, F). Consistent with our previous reporter profile, continuous PA predominantly induces trophectoderm differentiation (Figure 1E,F)^14^. By contrast, cells in PA-N differentiated to all three fates: a major fraction of cells downregulated expression of naïve epiblast markers, KLF4, KLF5, and TFCP2L1 and acquired expression of later epiblast markers, SFRP2, ETV4, and ETV5, indicative of early formative transition (Figure 1E,F, S1H). The remaining cells diverged along the extraembryonic fates: one merged towards the trophectoderm population differentiated in PA (Figure 1E,F); the second progressively upregulated hypoblast markers GATA6, PDGFRA, GATA4 and FOXA2 (Figure 1E,F).

### In vitro hypoblast differentiation resembles the emergence of nascent hypoblast in the embryo

We sought to compare the in vitro differentiation path with hypoblast formation in embryo development. We took advantage of a recent high-resolution embryo single-cell UMAP embedding^4^. This integrated embedding utilises entropy sort feature weighting (cESFW) for feature selection (3012 genes). The data covers pre-implantation (dpf 3 to 7) to in vitro cultured post-implantation (dpf 8 to 14) embryos (Figure 2A). The UMAP displays a continuous trajectory for formation of hypoblast (Figure 2B). Following the segregation of trophectoderm, ICM cells reach a branch point at dpf 5/6 expressing both ICM (NANOG, LAMA4) and early hypoblast (PDGFRA, GATA6) markers^5^. These cells resolve to either NANOG negative nascent hypoblast, or GATA6 negative nascent epiblast. The nascent hypoblast cells at dpf 6/7 begin to express FOXA2 and GATA4, two markers expressed exclusively in the hypoblast lineage in the pre-implantation blastocyst. The hypoblast continues towards hypoblast derivatives in extended culture samples (dpf 8-14 cells) (Figure 2A), with upregulation of APOB, PODXL, HABP2, which have been shown to be expressed in yolk-sac endoderm (Figure 2B) ^20,21^. The definitive endoderm, which emerges during post-implantation gastrulation stage embryogenesis, shares many markers, such as SOX17, GATA6, FOXA2, with the hypoblast lineage (Figure S2A). We therefore took advantage of a recent scRNA-seq resource of a ∼dpf19/cs7 gastrulation stage embryo^22^ to identify genes that are either lowly or non-expressed in definitive endoderm but are present in hypoblast, such as PDGFRA, RSPO3, HNF4A, LGALS2 (Figure 2B, S2B). These genes all show significant expression in nPSC-derived hypoblast (Figure S2C).

**Figure 2:**
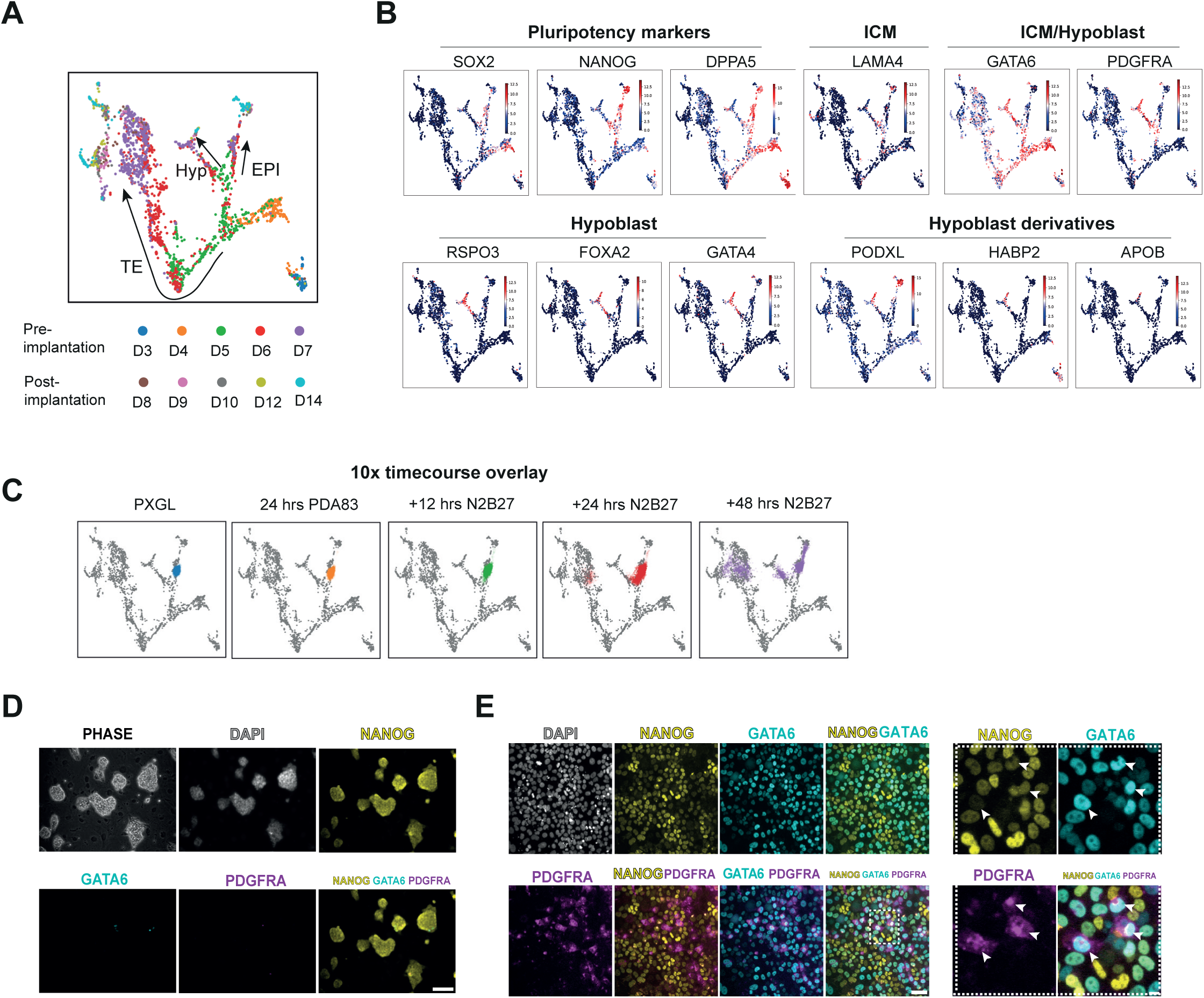
Comparative analysis of hypoblast differentiation with embryo lineage segregation trajectories. (A) UMAP embedding of integrated embryo single-cell datasets generated using entropy sorting-selected gene features as in Radley and Smith (2023). UMAP shows cells by embryo stage. (B) Log2 expression of pluripotency, hypoblast, and hypoblast derivative markers in (A), indicating lineage trajectories. (C) Projection of 10x scRNA-seq PA-N timecourse on embryo embedding using the same entropy sorting selected gene features as in (A). (D) Immunostaining for indicated markers on HNES1 GATA3:mKO2 cells cultured in PXGL on mouse embryonic feeders (MEFs), Scale Bar = 100 μM. (E) Immunostaining for indicated markers on HNES1 GATA3:mKO2 cells following one day of treatment with PD03+A83 and release into N2B27 for 16 hrs. Scale Bar = 50 μM. Zoomed-in inset scale bar = 5 μM

We projected the in vitro differentiation time course onto the embryo embedding based on transcriptome similarity over the 3012 gene set^4^ (Figure 2C). As expected, most naïve PSCs in PXGL position over dpf 6/7 naïve epiblast. The initial 24h of PDA83 treatment did not change this placement. Releasing into N2B27, however, saw a sub-population of cells shift to the epiblast/hypoblast branching point within 24 hours and reach the dpf 6/7 nascent blastocyst stage after 48 hours. In the embryo embedding, the nascent hypoblast emerges from a cluster of cells expressing NANOG, PDGFRA and GATA6. By immunofluorescence, we confirmed widespread co-expression of NANOG and GATA6 protein, as well as the presence of a subpopulation of triple-positive NANOG/GATA6/PDGFRA cells following release into N2B27 (Figure 2E). Moreover, NANOG/GATA6 double positive cells had low-mid expression of NANOG. These observations are consistent with previous descriptions of a mutually antagonistic interaction between these transcription factors in the mouse ICM^23,24^. Few cells cultured in PXGL co-express NANOG/GATA6 with no PDGFRA expression detected (Figure 2D). These results suggest that naïve PSC to hypoblast differentiation in PA-N proceeds via reversion to the epiblast/hypoblast branching stage of ICM.

### Naïve PSC to hypoblast differentiation requires FGF signalling

The preceding results show that MEK/ERK inhibition initially enables hypoblast differentiation potential of naïve PSCs, but the inhibition must be relieved for specification to occur. In the mouse ICM, the FGF/ERK/GATA6 signalling axis plays an instructive role in specifying hypoblast fate^25–29^. The FGF expressing cells activate FGF signalling in receptor-expressing cells, which leads to ERK-dependent upregulation of GATA6, downregulation of NANOG, and initiation of the hypoblast specification program. We surveyed FGF receptor and ligand expression in human embryo data and PA-N differentiation time course. In embryos, FGFR1 and FGFR2 are broadly expressed in ICM, epiblast, and hypoblast. Low expression of the FGF ligands, FGF4, was detected in ICM and followed by a prominent upregulation in nascent epiblast cells. This expression pattern resembles the restricted FGF4 expression in mouse ICM and epiblast^30,31^. Human naïve PSCs express FGF2 and FGF4 but lack an appreciable expression level of FGF receptors (Figure S3A). In vitro hypoblast differentiation is associated with the downregulation of FGF4/FGF2 ligands and upregulation of FGF receptors as well as GATA6 and PDGFRA upon release from PDA83 (Figure S3A).

We directly examined the function of FGF signalling during PA-N differentiation. We applied a FGF receptor inhibitor PD173074 (PD17) to inhibit the activation of FGF signalling and assayed hypoblast differentiation efficiency by OCT4-eGFP/SOX17-tdTomato reporter expression. We added PD17 either immediately following PDA83 induction (PA-PD17) or delayed for 24 hours after PDA83 (PA-N-PD17). We thus aimed to investigate FGF function at two hypoblast differentiation stages – during the epiblast/hypoblast branch point (PA-17) or after the branch point (PA-N-PD17) (Figure 3A). Flow cytometry revealed a dramatic reduction of SOX17-tdTomato + population from ∼14% to ∼2% in PA-PD17 condition (Figure 3B,C). However, delayed FGF inhibition in PA-N-PD17 condition failed to effectively block SOX17-tdTomato expression, as only a small reduction in SOX17-tdTomato expressing cells, from ∼14% to 11%, was recorded (Figure 3B,C), indicating that most cells released for 24 hrs in N2B27 had already been specified to hypoblast fate, and the establishment of FOXA2+ GATA4+ nascent hypoblast cells is not dependent on FGF signalling. These results point out that FGF signalling is crucial in initiating hypoblast lineage at the epiblast/hypoblast branch point but is not required for later progression.

**Figure 3:**
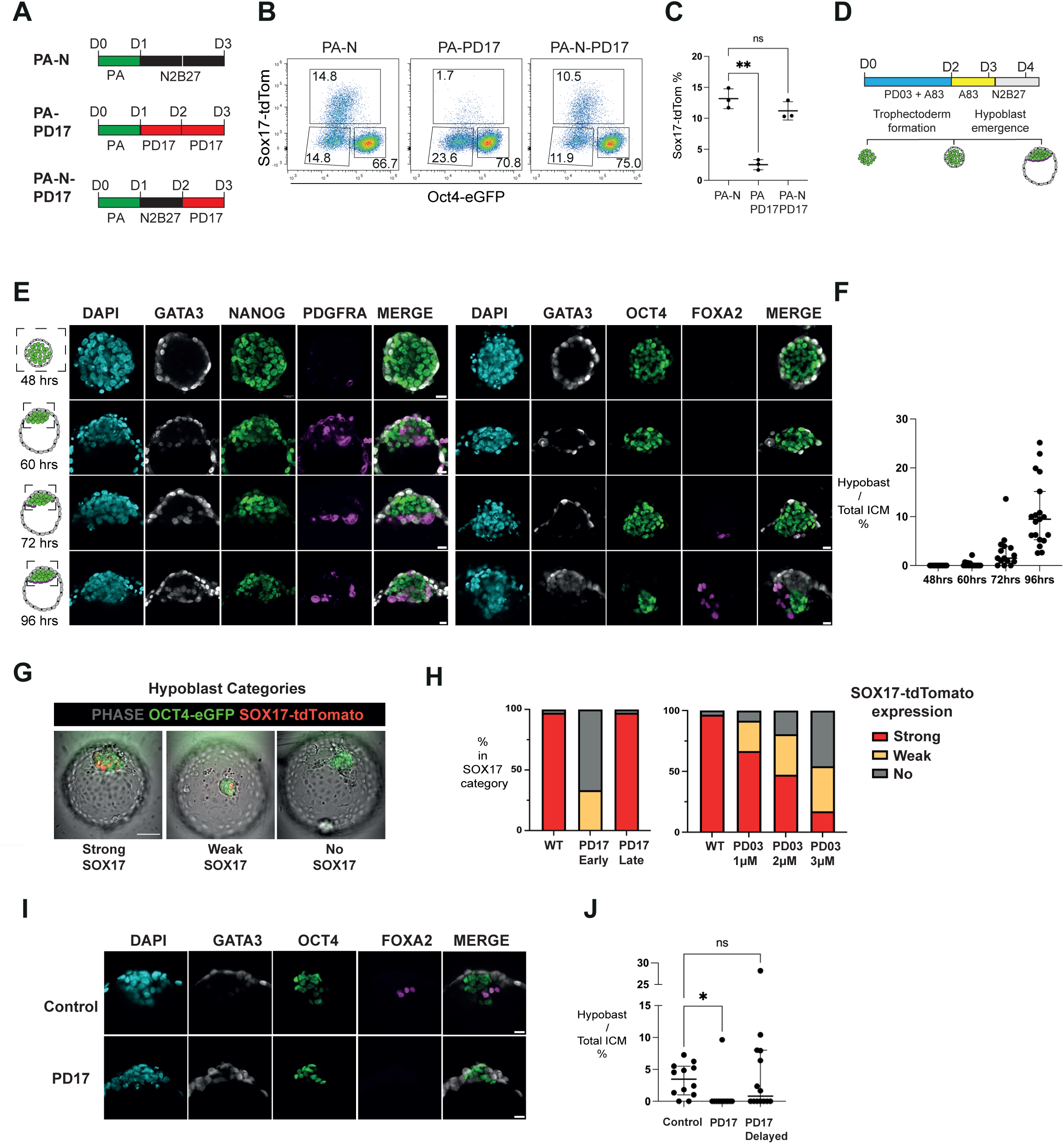
In vitro hypoblast differentiation requires FGF. (A) A schematic illustration of PA-N differentiation conditions with PD17 (0.5 μM) added at two subsequent time points following PDA83 (PA) treatment. (B) Flow cytometry analysis of OCT4-eGFP and SOX17-tdTomato expression following PA-N, PA-PD17, and PA-N-PD17 treatment. (C) Plot shows the SOX17-tdTomato-positive population quantified by flow cytometry analysis as in (B). Mean + S.D. from N=3 experiments. (D) A schematic illustration of blastoid formation protocol using PXGL naïve PSCs. (E) Immunostaining for indicated markers during blastoid formation using HNES1 GATA3:mkO2 cells. Scale Bar = 20 μM (F) Quantification of hypoblast (FOXA2+) versus ICM/Epiblast (OCT4+) cells at indicated time points during blastoid formation. Mean ratio + S.D. from N=3 experiments. (G) Live imaging of blastoids formed using niPSC-OS1 reporter cells, indicating three categories of the brightness of SOX17-tdTomato reporter. (H) Assay of SOX17-tdTomato expression in niPSC-OS1 blastoids formed with or without FGF signalling inhibitors. Inhibitors were added immediately following PDA83 treatment except for the PD17-delayed samples, where the PD17 was added 24 hours after release from PDA83. Bar chart shows the percentage of niPSC-OS1 blastoids according to the brightness of SOX17-tdTomato reporter expression as in (G). (I) Assay of hypoblast cells in HNES1 GATA3:mKO2 blastoids at 96 hrs with or without PD17. Images are immunostaining for indicated markers. Scale Bar = 20 μM. (J) Quantification of hypoblast (FOXA2+) versus ICM/Epiblast (OCT4+) cells in HNES1 GATA3:mKO2 blastoids at 96hr. PD17 was added either early or 24 hours delayed, as in (G). Mean ratio + S.D. from N=3 experiments.

### Hypoblast formation in blastoids also requires FGF/MEK/ERK signalling

We and others have previously reported the formation of a 3D blastocyst model, the blastoid, from naïve PSCs^16,17^. Immunostaining and single cell transcriptome analysis revealed that the fully expanded blastoid contains hypoblast cells^16,17^. However, the dynamics of hypoblast lineage segregation in the blastoid have not been described. We examined hypoblast lineage marker expression by immunofluorescence staining over the course of blastoid formation. Blastoids were formed by clustering naïve PSCs in non-adherent U-shaped wells in trophectoderm inductive PDA83 medium for 2 days, followed by one day of A83 treatment, and then one day of N2B27 alone (Figure 3D). We saw widespread upregulation of GATA6 after 48 hours in PDA83 (Figure S3B). Following medium exchange to A83, the early hypoblast marker PDGFRA appeared in some inner cells at 60 hrs and was widespread by 72 hours (Figure 3E). FOXA2 was detected later than PDGFRA, matching the sequence of activation observed in the blastocyst^5^. Moreover, FOXA2-positive cells were almost always positioned beneath the OCT4-positive epiblast population, indicating that this marker appears during or after spatial segregation of the hypoblast. The number of FOXA2+ cells was variable between blastoids. We quantified OCT4+ epiblast cells and FOXA2+ hypoblast cells in 15 blastoids at time points during blastoid formation. We saw sporadic FOXA2 in a minority of blastoids at 60 hours (Figure 3F). Over the following 2 days, the number of FOXA2 cells increased. By 96 hrs all blastoids examined contained FOXA2-positive cells, comprising from ∼2% to ∼25% of total inside cells.

We investigated whether FGF signalling underpins hypoblast formation in blastoids. We made blastoids from niPSC-OS1 and showed that inhibiting FGF signalling early in PD17, after release from PDA83, abolished nearly all SOX17-tdTomato+ cells in blastoids examined at 96 hrs (Figure 3G,H, S4B). 24 hours of delay in application of PD17 showed a reduced effect. 35 of 36 blastoids contained bright SOX17-tdTomato+ cells (Figure 3G,H, S4A). Likewise, we made blastoids using HNES1 GATA3:mkO2 cells and similarly showed that inhibiting FGF signalling directly after release from PDA83, abolished nearly all FOXA2+ hypoblast cells in blastoids (Figure 3I, J). Delayed application of PD17 led to a mild reduction in FOXA2+ (Figure 3I, J). In mouse, FGF promotes hypoblast specification via activation of MEK/ERK signaling^32^. We examined hypoblast formation in niPSC-OS1 blastoids subjected to MEK/ERK inhibition with PD03 following release from PDA83. We found that 2-3 uM PD03 impeded Sox17-tdTomato+expression effectively (Figure 3G,H, S4B). However, lower concentrations of PD03 (1uM) showed only partial inhibition (Figure 3G,H, S4A). Taken together, these results indicate that FGF/MEK/ERK signalling is required in hypoblast specification in human blastoids.

### Signalling environment enhances hypoblast differentiation

Hypoblast-like cells constitute less than 20% of cells after 2D differentiation in PA-N. We investigated whether this could be increased by adjusting the signalling environment. Release from 24hrs PDA83 treatment into varying concentrations of FGF2 (up to 250 ng/ml) or FGF4 led to only a modest increase in the number of SOX17-tdTomato + cells (Figure S5A,B,C), indicating that autocrine FGF production is near saturating. Supplementation with Activin or the GSK3 inhibitor, Chiron 99021, two components purported to promote mouse and human extraembryonic endoderm differentiation^20,33,34^, reduced the SOX17-tdTomato population (Figure S5D,E).

The reporter assay and differentiation trajectory analysis showed that epiblast and trophectoderm cells are major populations in the PA-N differentiation. We reasoned that hypoblast differentiation may be enhanced by reducing the epiblast and trophectoderm potential. We tested Activin/Nodal inhibition by A83-01 to destabilise pluripotency^35^ and tankyrase inhibition by XAV-939 to limit trophectoderm^15^. The addition of A83-01 reduced OCT4 positive population by flow cytometry analysis (Figure S5F), and the addition of XAV-939 reduced the trophectoderm (SOX17^-^/OCT4^-^) population as expected (Figure S5F). However, treatment with single inhibition did not dramatically increase SOX17+ population. Either combined with FGF2 (25 ng/ml) synergistically increased the SOX17-tdTomato+ population (Figure 4B). Combining all three inhibitors FGF2+A8301+XAV939 (hereby referred to as FA83X) resulted in the most significant increase in SOX17-tdTomato+ population (Figure 4B, C). One day of PDA83 treatment followed by two days of FA83X culture yields ∼40% SOX17+ cells (Figure 4B, C). We sorted SOX17+ cells by flow cytometry and examined the gene expression profile of the FA83X differentiation time course by bulk RNA sequencing analysis (Figure S5G). We saw a sequential upregulation of early and late hypoblast signature genes after 24 hours and 48 hours in FA83X (Figure S5G).

**Figure 4:**
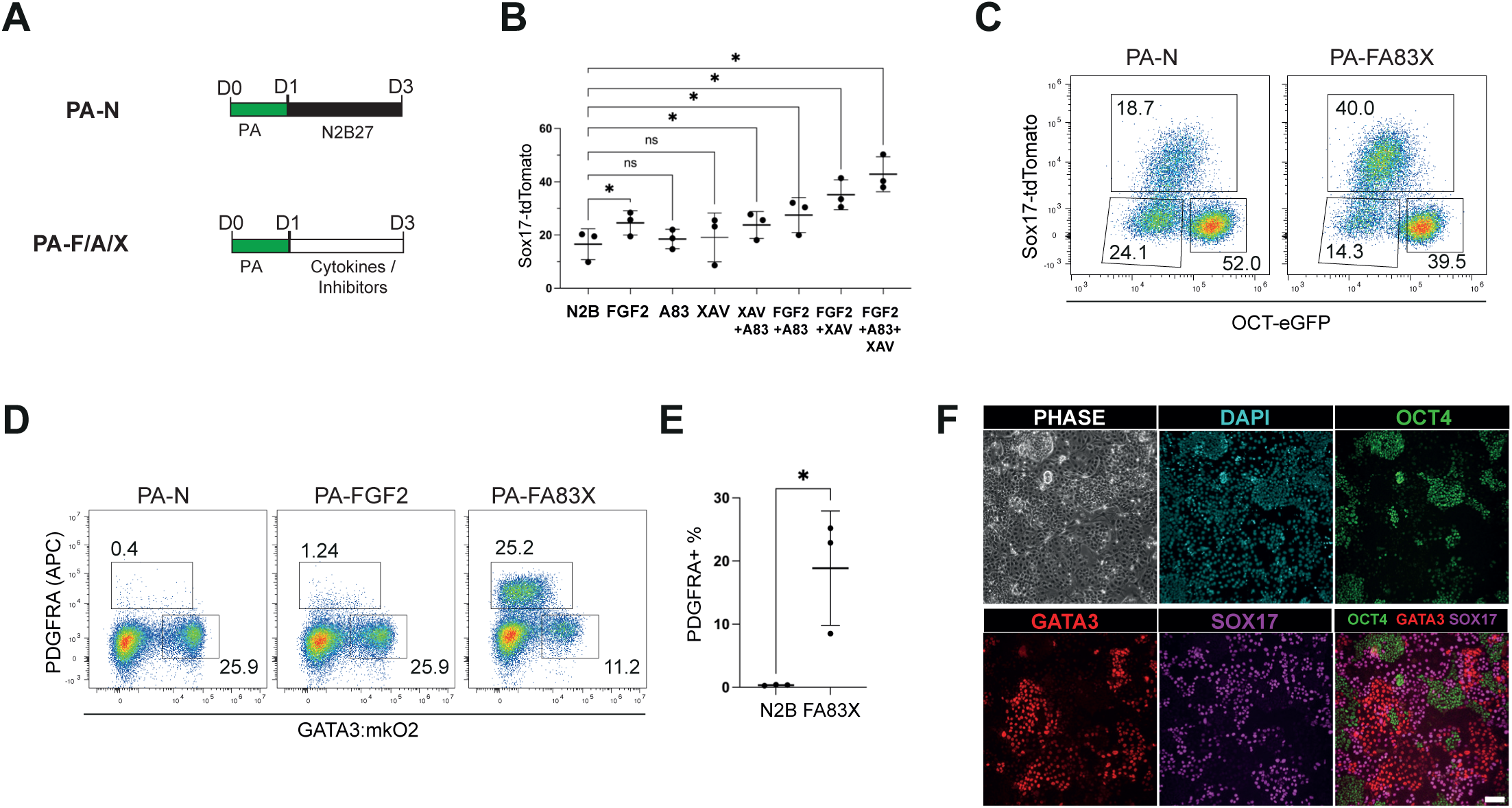
Signalling influence on hypoblast differentiation. (A) Schematic of differentiation conditions for testing of small molecule inhibitors or cytokines that improve hypoblast efficiency compared to PA-N. (B) The percentage of SOX17-tdTomato positive population by flow cytometry analysis in niPSC-OS1 culture after differentiation with indicated testing factors added singly or combined. Mean + S.D. from N=3 experiments. (C) Flow cytometry analysis shows subpopulations of OCT4-eGFP+, SOX17-tdTomato+, and double-negative populations following three days of differentiation in PA-N or PA-FA83X. (D) Flow cytometry analysis of PDGFRA and GATA3:mKO2 expression in HNES1GATA3:mkO2 culture following three days of differentiation in indicated conditions. (E) Percentage of PDGFRA-APC-positive population by flow cytometry as in (D). Mean + S.D. from N=3 experiments. (F) Immunostaining for indicated markers for HNES1 GATA3:mkO2 following differentiation in PA-FA83X (3 days). Scale Bar = 100 μM.

We examined hypoblast differentiation in HNES1 GATA3:mKO2 reporter cell lines by staining for PDGFRA protein. This cell line differentiated very efficiently to trophectoderm and generated less than 1% PDGFRA+ cells in PA-N^14^(Figure 4D). Adding FGF2 marginally increased the PDGFRA+ population, but FA83X yielded ∼25% PDGFRA+ cells (Figure 4D, E). Immunofluorescence staining confirmed the expression of hypoblast markers SOX17 and FOXA2 in the PDGFRA+ population (Figure 4F, S5H). These results confirmed the effect of FA83X in 2D adherent hypoblast differentiation and supported the hypothesis that FGF-instructed hypoblast differentiation maybe enhanced further by reducing epiblast and trophectoderm.

We then tested FA83X on hypoblast differentiation in blastoids made from HNES1:mKO2 reporter cells. We added FA83X following 48 hours of PDA83 for two days. Blastocyst-like morphology with GATA3+ trophectoderm layer and the blastocoel-like cavities initially formed by 72 hours, however, the cavities subsequently collapsed between 72 and 96, likely due to inhibition of YAP downstream of XAV939 and impairment of trophoblast differentiation^15,36^. Nevertheless, we observed a dramatic increase of FOXA2+ cells at 96 hours. The resultant clusters generally formed an organised bilayer of hypoblast underlying a large epiblast population (Figure S5 I, J, K).

### PA-FA83X differentiation follows the embryo hypoblast development trajectory

We performed 10x single-cell RNA sequencing on the PA-FA83X time course and produced an integrated UMAP embedding for PA-N and PA-FA83X differentiation. PA-N and PA-FA83X cells largely intermingled along the time course (Figure 5A). Leiden cluster analysis identified 10 clusters (Figure 5B). Marker expression assigned three main trajectories along the differentiation time course. Naïve PSCs (cluster 0) either transitioned to formative epiblast (clusters 1, 8, 9, 10) or reached a branching population (cluster 2) from which they diverged to hypoblast-like (clusters 3, 4) or trophectoderm-like fate (cluster 6, 7) (Figure 5B). By the end of the differentiation, a distinct cluster (Cluster 5) is present. This cluster showed enriched expression in genes associated with extraembryonic mesoderm in marmoset peri-implantation embryos, including genes related to the epithelial-to-mesenchymal transition (BMP4, LEF1, ZEB2, ETS1, MEIS2, FLI1, and HAND1) and extracellular matrix organisation (COL4A2, ITGA3, HAPLN1, KDR)^21^ (Figure 5C, Figure S6A). We therefore annotated this cluster as ExEM-like cells.

**Figure 5:**
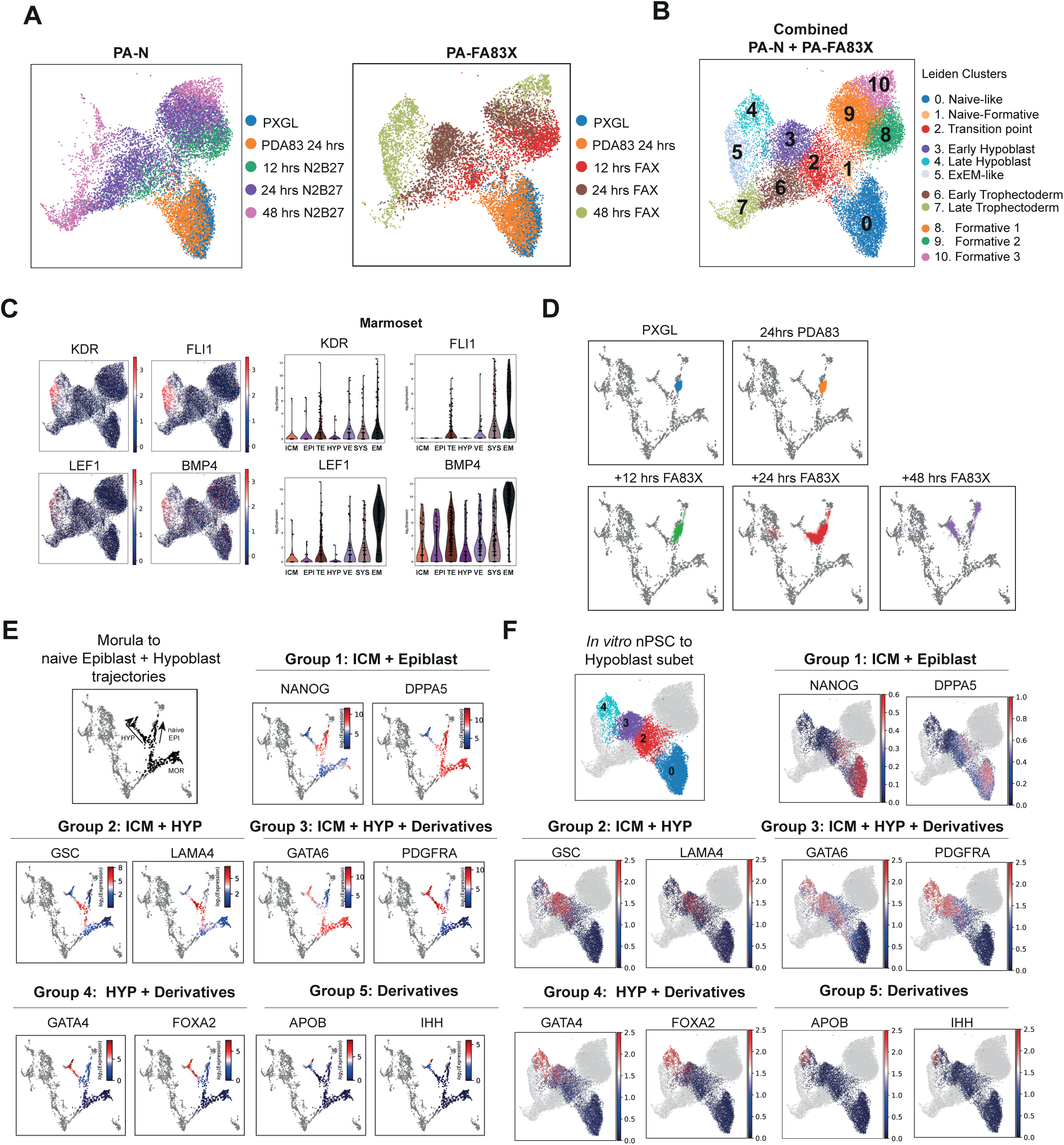
Single-cell sequencing analysis of the PA-FA83X differentiation. (A) UMAP of PA-N and PA-FA83X timecourses. (B) Combined PA-N and PA-FA83X UMAPs, coloured according to identified Leiden clusters. (C) Log2 expression of selected extra-embryonic mesoderm markers in (A). Markers are upregulated in extra-embryonic mesoderm in marmoset early embryo single-cell dataset (Bergmann 2022 ^21^). (D) Projection of 10x scRNA-seq PA-FA83X timecourse on embryo embedding. (E) Smoothed expression of indicated genes in embryo UMAP embedding subsetted to morula (D4), ICM (D5), naïve epiblast (D6+D7), and hypoblast cells (D6-D14). (F) Expression of indicated genes in combined PA-N and PA-FA83X UMAP subsetted to naïve epiblast, transitional ICM, early hypoblast, and late hypoblast.

We focused on characterisation of hypoblast formation in PA-FA83X. We first examined global transcriptome similarity to embryo lineages and projected the PA-FA83X time course on the integrated embryo UMAP embedding (Figure 5D). Like the PA-N time course, PA-FA83X differentiation begins with a transition towards a branching point where ICM cells bifurcate to epiblast and hypoblast before progressing to dpf 6/7 nascent hypoblast. Compared to PA-N, FA83X accelerated hypoblast lineage maturation. After 48 hours in FA83X, cells are positioned adjacent to hypoblast derivatives while PA-N cells mostly remain over nascent hypoblast.

We collated panels of genes enriched in epiblast and hypoblast lineages and examined their expression dynamics along hypoblast development in the embryo embedding and in the FA83X differentiation time course UMAP (cluster 0, 2, 3, 4) (Figure 5E,F). The genes are grouped into 5 classes according to expression patterns (Figure 5E). Group 1 are pluripotent genes, including NANOG and DPPA5, which are expressed in ICM and sustained in the epiblast/hypoblast branch and epiblast but switched off in the nascent hypoblast. Group 2 are ICM markers and mostly sustained in epiblast/hypoblast branch, but downregulated in the epiblast and nascent hypoblast, such as GSC and LAMA4. Group 3 are expressed in ICM and throughout hypoblast development (GATA6, PDGFRA). Group 4 and Group 5 are hypoblast-specific markers that appear either in mature blastocyst hypoblast (FOXA2, GATA4) or late in hypoblast derivatives (APOB, IHH). In vitro hypoblast differentiation exhibited similar gene expression dynamics to the trajectory in the real embryo from the ICM-like branching population (Figure 5F). In vitro differentiation began with NANOG and DAPPA5 expressing epiblast-like cells (cluster 0), Cells expressed ICM and early hypoblast markers including GATA6, PDGFRA, GSC and LAMA4 within 12 hrs in FA83X after release from PDA83 (mostly cluster 2). Over the following 36 hours in FA83X the expression of epiblast and ICM markers NANOG, DPPA5, GSC and LAMA4 gradually diminished, and hypoblast-specific gene expression was up-regulated (cluster 3,4). After 48 hours of differentiation in FA83X, a group of peri/post-implantation yolk-sac endoderm markers are detected including IHH and APOB, two markers expressed in primate hypoblast lineages^21^ (Figure S2B). By pseudo-temporal ordering, we delineated the sequential activation of a spectrum of markers along the morula to hypoblast trajectory in the embryo embedding (Figure S6B). We plotted these markers in our 10x in vitro timecourse and found they broadly mirrored the temporal upregulation of the hypoblast markers in embryos (Figure S6C).

Taken together, the global transcriptome and marker expression comparison between embryo and in vitro differentiation support the conclusion that both PA-N and PA-FA83X differentiation reflect hypoblast lineage specification dynamics in the blastocyst. In vitro differentiation is initiated with the onset of co-expression of ICM and early hypoblast markers, partially overlapping with the late ICM and epiblast/hypoblast branch in the embryo embedding (Figure 2C, 5D). We refer to this as a naïve epiblast towards ICM-like reversion, a pivotal step in enabling hypoblast lineage specification in vitro.

### Hypoblast lineage specification in the human embryo depends on FGF signalling

These stem-cell based studies indicate that FGF is critical to initiate hypoblast differentiation from an ICM-like state. We therefore examined whether FGF signalling plays a similar role in the embryo. To capture the window of emerging hypoblast, we applied PD17 to early day 5 blastocysts, a stage when most ICM cells are positioned before the epiblast/hypoblast branch point (Figure 2A). We removed the zona pellucida to ensure free access of the inhibitor to the embryo. The freshly thawed dpf 5 embryos were cultured either in N2B27 basal medium or with 0.5 μM PD17 for 24-40 hours until they developed fully expanded blastocoel cavities with a recognisable ICM and the appearance of smaller trophectoderm cells, traits of late dpf 6/7 blastocysts. The embryos were fixed and immunostained for NANOG (epiblast), SOX17 and FOXA2 (hypoblast), and GATA3 (trophectoderm) (Figure 6A). We examined thirteen PD17-treated embryos and eleven control (N2B27) embryos in three independent experiments. All control embryos contained SOX17 or FOXA2-positive hypoblast cells within the ICM, exclusive of NANOG-positive cells. In PD17-treated groups, two embryos did not contain an ICM. Seven out of the remaining eleven embryos contained NANOG positive epiblast but were devoid of hypoblast (Figure 6B,C). PD17-treated embryos contained on average a higher number of epiblast cells compared to the control embryos, which is consistent with a hypoblast to epiblast lineage switch (Figure 6D). This result mirrors the phenotype of mouse embryos treated with FGFR or MEK inhibitors^28^. The PD17-treated embryos had an intact blastocyst cavity, indicating no general cellular toxicity. However, four of the 11 PD17-treated embryos escaped the hypoblast lineage blockage, displaying a similar hypoblast cell number as the control condition (Figure 6B). The developmental pace of human IVF embryos varies and staging based on morphology is not precise. We speculate that the embryos that formed hypoblast in PD17 may have been at a more advanced stage when exposed to PD17.

**Figure 6:**
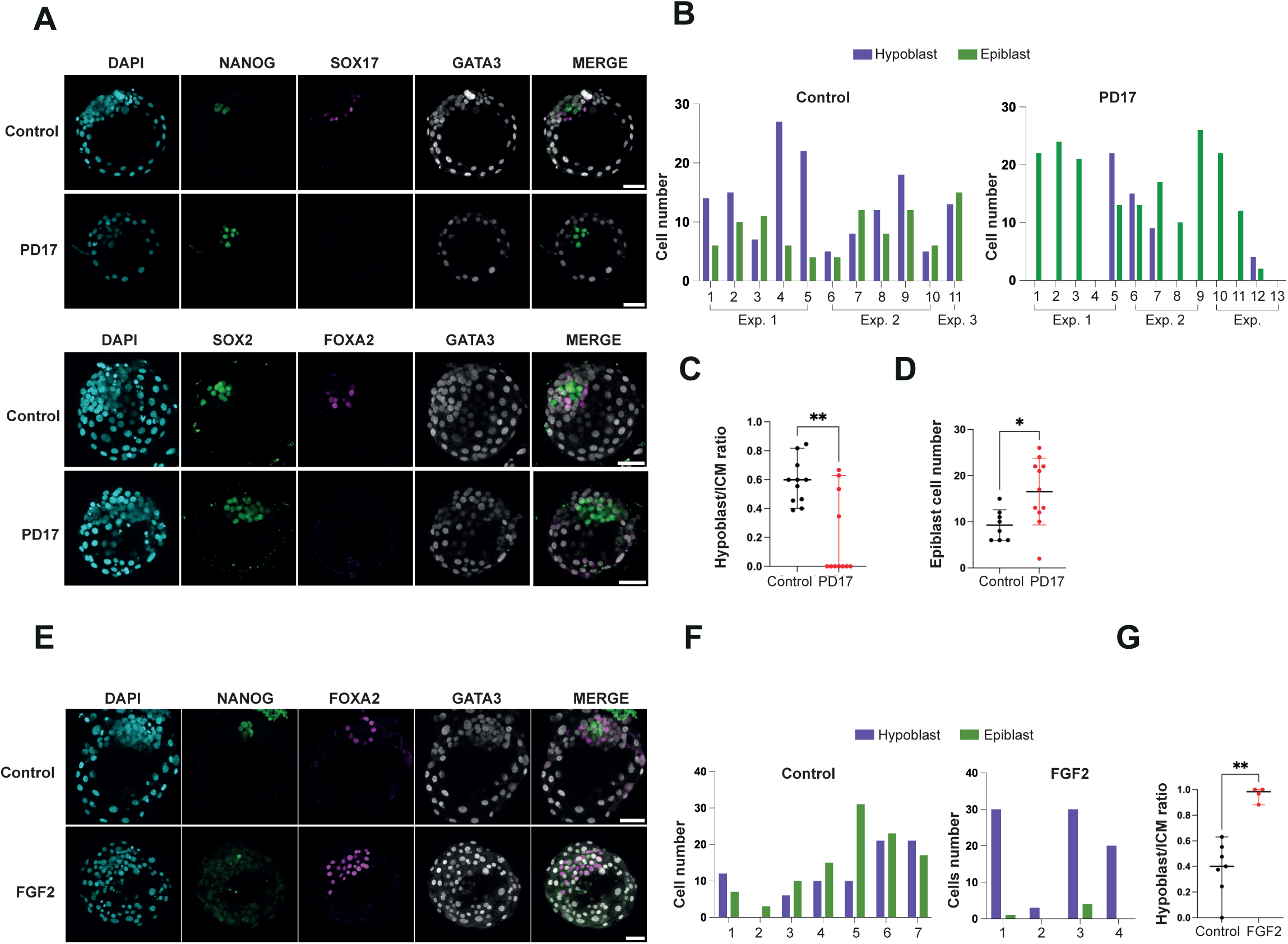
FGF signalling drives human hypoblast in blastocyst. (A) Immunostaining for indicated markers in fully expanded blastocyst following 24 – 44 hours culture either in N2B27 (control) or N2B27 containing PD17 (0.5 μM) after thawing of vitrified human embryos at dpf 5. Scale Bar = 50 μM (B) Quantification of epiblast (SOX2+ or NANOG+) and hypoblast (SOX17+ or FOXA2+) cell numbers in control and PD17-treated embryos in three experiment. X-axis indicates each embryo assayed. (C) Hypoblast to total ICM (epiblast & hypoblast) ratio for control and PD17 treated embryos. (D) Comparison of epiblast cell numbers of control and PD17 treated embryos. (E) Immunostaining for indicated markers in fully expanded blastocyst after 24 – 44 hours culture in N2B27 or FGF2 (250ng/mL) following thawing of vitrified D5 human embryos. Scale Bar = 50 μM (F) Quantification of epiblast and hypoblast numbers in control and FGF2 treated embryos. (G) Hypoblast to total ICM (epiblast & hypoblast) ratio for control and FGF2 treated embryos.

Finally, to examine whether FGF can promote hypoblast formation, we cultured dpf 5 embryos until the late blastocyst stage in medium supplemented with FGF2(250 ng/ml). Quantification of FOXA2 and NANOG expressing cells within the ICM showed a dramatic increase in hypoblast cell number at the expense of epiblast in 4 out of 4 FGF-treated embryos (Figure 6E,F, G). These results together reveal the critical role of FGF in hypoblast lineage specification in early human blastocysts and establish consistency with findings in the mouse embryo^28^.

## DISCUSSION

Embryogenesis is a process of dynamic cell fate specification and spatial organisation, which must be well-coordinated to ensure developmental robustness. The limited research access to early embryos has hampered studies of human embryogenesis. Stem cell-based embryo models are, therefore promising as a scalable alternative. However, to be a useful model for studying the regulatory mechanisms of cell fate specification during development a model must faithfully capture the developmental dynamics and signalling logic in the early embryo. In this study, we establish an in vitro cellular model for human hypoblast lineage segregation. Following a short period of exposure to MEK/ERK and Activin/Nodal inhibition, human naïve pluripotent stem cells transition to an ICM-like state whence they give rise to hypoblast and trophectoderm. Single-cell transcriptome analysis indicates that the in vitro ICM-like state to hypoblast differentiation mirrors the ICM to hypoblast trajectory in the embryo. Authentic nascent hypoblast emerges in in vitro cultures that closely approximates dpf 6/7 hypoblast *in vivo* and further develops features of yolk-sac-like endoderm.

A key finding in hypoblast differentiation in both adherent and blastoid cultures is the indispensable role of FGF signalling. Our experiments have uncovered a transient developmental window during which FGF signalling plays a critical role in determining the fate of the hypoblast. This 24-hour window in the differentiation timeline aligns with the early dpf5 embryo before the divergence of hypoblast and epiblast within ICMs. Our in vitro findings align with *in vivo* results, as inhibiting FGF signalling in dpf 5 embryos led to the suppression of hypoblast formation. Additionally, we demonstrated that the addition of FGF2 to dpf 5 embryos resulted in the conversion of nearly all ICM cells into hypoblast cells, further substantiating the pivotal role of this signalling pathway in hypoblast specification. It must be noted that two previous studies with human embryos failed to establish a requirement for FGF signalling in hypoblast lineage specification^37,38^. However, those experiments used the MEK inhibitor PD0325901 at concentrations of only 0.5 μM ^31^ or 1.0 μM ^32^ while we found a concentration of 2.0 μM is required to suppress hypoblast specification in blastoids. Thus, by enabling more systematic investigation, the stem cell models have clarified the signalling mechanism directing hypoblast specification in human embryo.

## LIMITATIONS OF THE STUDY

Currently there is limited transcriptome data available on post-implantation stage human embryo hypoblast derivatives. Consequently, we cannot accurately distinguish the various derivatives of the hypoblast generated by in vitro differentiation. Moreover, additional single cell transcriptome data for dpf 5-dpf 6 embryos may provide greater resolution of, and more markers for, the late ICM hypoblast-epiblast branch point. The limited availability of human embryos and the challenges in precisely determining their developmental stage constrain the scale and scope of our study. In addition, we no longer have access to morula stage embryos and therefore cannot repeat inhibitor experiments exactly as performed in earlier studies. Two factors that may have influenced the outcome of previous studies, the half-life of inhibitors in culture with embryos and the effect of the zona pellucida on access to the embryo, have not been quantified.

## ACKNOWLEDGEMENTS

We thank Austin Smith for critical comments and for editing the manuscript. Somayyeh Sadat Tahajjodi provided laboratory assistance. We also acknowledge Corin Liddle and Exeter Bioscience Imaging Centre for confocal imaging. We thank Paul O’Neill and Jemima Onime from Exeter Sequencing Facility for their dedicated support in data analysis. This research project was funded by Biotechnology and Biological Sciences Research Council (BBSRC) with a research grant (BB/V017128/1). This project utilised equipment funded by the Wellcome Trust Institutional Strategic Support Fund (WT097835MF), Wellcome Trust Multi-User Equipment Awards (WT101650MA, 218247/Z/19/Z) and BBSRC LOLA award (BB/K003240/1) and Medical Research Council (MRC) Clinical Infrastructure Funding (MR/M008924/1). PWZ is the Canada Research Chair in Stem Cell Bioengineering. PWZ receives funding from NSERC Discovery Grant (RGPIN-2020-06496) and CIHR Foundation Grant (FDN 154283). SL is supported by a Wellcome GW4-CAT fellowship. (222850/Z/21/Z).

## AUTHOR CONTRIBUTIONS

Conceptualisation, GG, AD; Investigation, AD, ECS, YT; Analysis, AD, AR, TH; Methodology, FC, SL; Writing, GG, AD; Supervision GG, JN, PZ, JM

## DECLARATION OF INTERESTS

GG is an inventor on a patent relating to human naïve pluripotent stem cells filed by the University of Cambridge. SL reports non-financial support from Pfizer outside the submitted work.

## MATERIALS AND METHODS

### Thawing and culture of human embryos

Use of supernumerary embryos is approved by the Multi-Centre Research Ethics Committee, approval 04/MRE03/44, Integrated Research Application System (IRAS) 21/PR/1231 and licensed by the Human Embryology and Fertilization Authority of the United Kingdom under the research license R0178.

Surplus frozen human embryos were donated for research after informed consent by couples following in vitro fertilization treatment. Early D5 blastocysts were thawed following specific IVF clinic instructions and cultured in equilibrated N2B27 medium under mineral oil in a humidified incubator at 37 degrees, 7% CO_2_ and 5% O_2_ for 1-2h before assessing viability and suitability for subsequent treatment.

### Human embryo treatment

The zona pellucida was removed from early D5 blastocysts (unless otherwise stated) using acid Tyrode solution (Gibco) prior to start of the treatment. Human early Day 5 embryos were treated with either: 0.5µM PD17, 250ng/ml FGF2 (all in N2B27) or maintained in N2B27 (+DMSO) as control. Blastocysts were cultured in NUNC IVF dishes (#150260, Thermo Fisher) in a humidified incubator at 37 degrees, 7% CO2 and 5% O2 for the duration of the treatment.

### Immunostaining of human embryos

The treatment was maintained until the embryos reached the late D6/D7 stage (30-48h), assessed by blastocoele size^5^, when embryos were fixed in 4% paraformaldehyde in PBS at room temperature for 15 min. Blastocysts were then rinsed in 3mg/ml polyvinylpyrrolidone (Sigma-Aldrich) (PBS/PVP) prior to permeabilization in 0.25% Triton X-100 (Sigma-Aldrich) for 30 min and blocking in 0.1% bovine serum albumin, 0.01% Tween 20 (Sigma-Aldrich) and 2% donkey serum for 2-3 hours at room temperature. Primary and secondary antibodies were diluted in blocking buffer. Primary antibody incubation was performed overnight at 4 degrees, followed by three rinses of 15 min in blocking buffer. Secondary antibody incubation was performed at room temperature, in the dark, for 1-2h followed by three additional rinses in blocking buffer for 15 min. Blastocyst were imaged through a poly-D-lysine-coated Mattek dish (P356-0-14) in blocking buffer.

### Imaging and analysis of immunostained human embryos

Stained human embryos were imaged with Leica Stellaris 8 confocal microscope, objective: HC PL APO CS2 40x/1.10 water. Confocal images were analysed using FIJI. The number of epiblast or hypoblast cells were counted using the “Cell Counter” plugin in FIJI.

### Human naïve PSC culture

Embryo-derived human naïve pluripotent cells (HNES1) or reprogrammed naïve pluripotent cells (nIPSCs) were propagated in PXGL medium as described in^10^. PXGL comprises 1 μM PD0325901 (P), 2 μM XAV939 (X), 2 μM Gö6983 (G) and 10 ng/mL human LIF (L) in N2B27 medium. Cells were cultured on mitomycin-inactivated MEF feeders. Rho associated kinase inhibitor (10μM, Y-27632) and Geltrex (0.5 μl/cm^2^ well surface area, Thermo Fisher Scientific, A1413302) were added to media during replating. Cells were passaged by dissociation with Accutase (Biolegend, 423201) every 3-4 days.

### Differentiation

Human naïve cells were plated in PXGL with Y-27632 (10μM) on Geltrex coated 6 well plates. The next day, cultures were washed with PBS and medium was exchanged to N2B27 containing chemical inhibitors and cytokines. Medium was refreshed or exchanged every day until the end of the assay. Concentrations of small molecules/cytokines were optimised for the cell line being assayed. For PDA83 treatment, PD03: 1 or 2 μM; A83-01: 1 or 2 μM. For FA83X hypoblast induction: FGF2: 25 ng/ml; A83-01: 1 μM; XAV: 2 μM.

### CRISPR-mediated generation of OCT4-GFP/SOX17-tdTomato nIPSCs

nIPSCs reprogrammed from human dermal fibroblasts (HDF16) were transfected with 3 μg targeting vector pUC19-OCT4-T2A-NLS-EmGFP-P2A-Puro (kind gift from Timo Otonkoski; Addgene plasmid #89992) and 3 μg of PX459 plasmid (kind gift from Feng Zhang; Addgene plasmid ##62988) expressing both cas9 and a cloned gRNA (5’-TCTCCCATGCATTCAAACTG-3’) to target near to the stop codon of OCT4. Correctly targeted colonies were selected by Puromycin selection for 3 days and checked for GFP expression. OCT4-T2A-GFP nIPSCs were then expanded and transfected with 3 μg SOX17-H2B-tdTomato-PGK-Neo targeting vector (kind gift from Ali H. Brivanlou) and 3 μg of PX459 plasmid expressing cas9 and a cloned gRNA (5’-CAACTATCCTGACGTGTGAC-3’) to target the stop codon near SOX17. Continuous selection with G418 for two passages purified potentially targeted cells. At least 40 colonies were picked, expanded, and screened for SOX17-tdTomato expression using 1 day pulse of PDA83 followed by release into N2B27. Correctly targeted OCT4-GFP/SOX17-tdTomato colonies were validated by differentiation in trophectoderm-inductive conditions to assay for disappearance of OCT4-GFP signal, and co-staining of SOX17-tdTomao with SOX17 antibody in hypoblast inductive conditions.

### Generation of blastoids

PXGL cultures of naive stem cells in exponential growth were dissociated with TrypLE for 5 min and spun in wash buffer (DMEM/F12 supplemented with 0.1% BSA). The cell pellet was resuspended in PD+A83+Y medium. For HNES1 GATA3:mKO2 cell lines we used 2 μM PD0325901, 2 μM A83-01 and 10 μM Y-27632 resuspended in N2B27 media. For niPSC cell lines we added 0.5 μM LPA to PD+A83+Y to increase cavitation efficiency. Cell numbers were counted using a haemocytometer. We added ∼20K cells to 10 ml of PD+A83+Y media. 50 μl of the cell suspension was dispensed to each well of an ultra-low attachment multiple-well plate (Cell Carrier Spheroid ULA 96-well Microplates, 6055330) with a multichannel pipette thereby aiming for ∼100cells/well. Plates were centrifuged at 1400 RPM for 5 min at room temperature to cluster cells at the bottom of the wells. After 48 h, the media in the well was changed to pre-warmed N2B27 supplemented with 0.5 μM A83-01 by careful aspiration using a multi-channel pipette. At the end of day 3, media was changed to N2B27 medium devoid of all inhibitors. Blastoids fixed at 96 hrs with 4% formaldehyde and transferred to PBS for storage at 4c.

### Immunostaining of blastoids

Fixed blastoids stored in PBS were permeabilized with 0.3% Triton X-100 in PBS (0.3% PBSTx) for 15 min. Blocking was performed in blocking buffer (1%BSA in 0.1% PBSTx) at room temperature for 1 hr. Primary antibodies were diluted in blocking buffer and incubated with blastoids for at least 16 hours overnight at 4c. Blastoids were washed with blocking buffer 4 times, before incubation of fluorophore-conjugated secondary antibody (1:1000 diluted in blocking buffer) for 1 hour at room temperature. Blastoids were washed three times with blocking buffer and followed by one wash in in PBS before addition of DAPI (500 ng/mL) in PBS and incubated for 15 mins. Stained blastoids were rinsed with PBS and were stored for imaging.

### Imaging of blastoids

Stained blastoids were imaged with Airyscan LSM880 with a D LCI Plan-Apochromat 25x/0.8 Imm Corr DIC M27 objective with oil immersion. Images were acquired with Airyscan FAST Mode for ICM cell quantification or with R-S mode of representative high-resolution images. Confocal images were analysed in IMARIS, and cells classified using the automated spot detection module, manually checked, and quantified.

### Transcriptome Sequencing

For 10x Genomics Chromium Single Cell sequencing, samples were multiplexed using 3’ CellPlex Kit Set A. Single cell capture and library preparation was performed on a Chromium Controller using the Chromium GEM Single Cell 3’ Kit version 3.1 as per the user guide, targeting a recovery of 3,000 cells per sample. We targeted a gene expression (GEX) coverage of 40,000-50,000 reads per cell, and 10,000-12,0000 reads per cell multiplex oligo. Sequencing was performed on a Novaseq 6000 instrument generating paired end reads (28×10×10×90 bases). Sample demultiplexing, alignment, and quantification of barcode counts was performed using CellRanger Multi.

Scanpy was used to read and analyse raw read counts from the CellRanger output. Cells expressing fewer than 15,000 counts, more than 100,0000 counts, or more than 15% mitochondrial reads were filtered out. The resultant count matrix was normalised and log-transformed using reciple_zheng_17. The top 1000 highly variable genes were identified for initial dimensionality reduction with PCA prior to non-linear dimensionality reduction using UMAP. Cells visualised in UMAP plots were coloured according to individual marker gene expression values, and the Leiden algorithm (resolution 0.8) was used to identify cell clusters.

### Comparison of in vitro cultures against human embryo reference

To relate samples from our in vitro sequencing data to the Radley et al. 2023 human embryo embedding^4^, we took the UMAP model object from their paper and used the *umap.transform* function to position each individual sample from our data into the UMAP latent space. In doing so, we can identify which samples in the early human embryo our in vitro samples are most transcriptionally similar to, according to an unbiased set of 3012 genes.

### Hypoblast gene activation pseudotime ordering

We took the UMAP embedding defined by Radley et al. (2023)^4^ and focussed on the cells stages comprising morula, ICM, Epi/Hyp branch point, epiblast and hypoblast. Pseudotime along the hypoblast branch was calculated using the Slingshot R package^39^. Gene expression values were smoothed by taking the average gene expression for each cell and its 30 most similar cells according to the 3012 highly structured genes identified by Radley et al. (2023)^4^. Logistic regression curves were fit to the smoothed expression profiles versus the hypoblast pseudotime. The hypoblast pseudotime value that corresponds to the half-way point on the *y*-axis of the logistic curve indicates the tipping point between a gene being active versus inactive.

## Notes

### Competing Interest Statement

GG is an inventor on a patent relating to human naive pluripotent stem cells filed by the University of Cambridge.

