## supplement figures for "Human naïve stem cell models reveal the role of FGF in hypoblast specification in the human embryo"

### Supplemental Figure 1

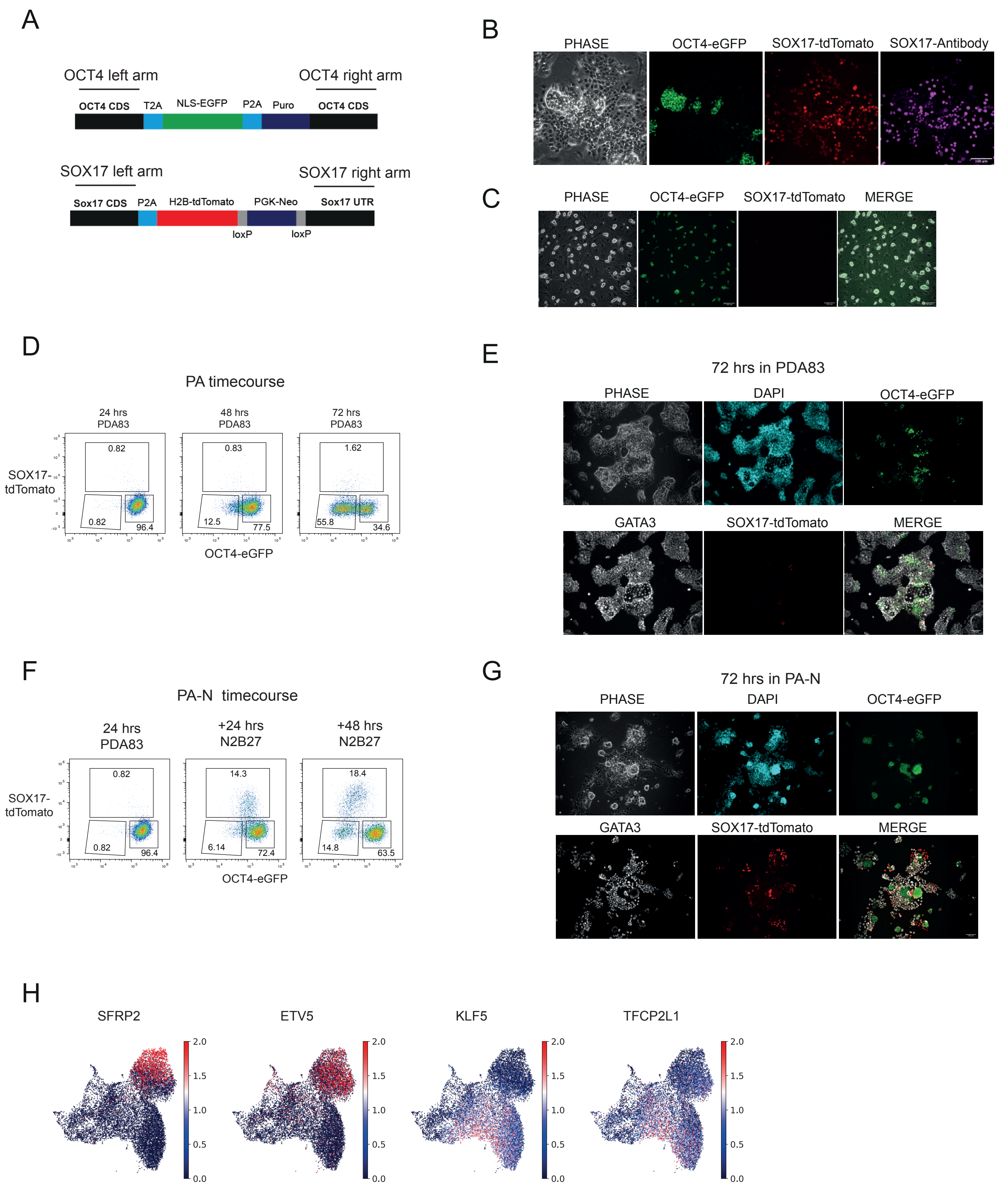

### Supplemental Figure 2

## A

##### Shared Hypoblast and Definitive endoderm

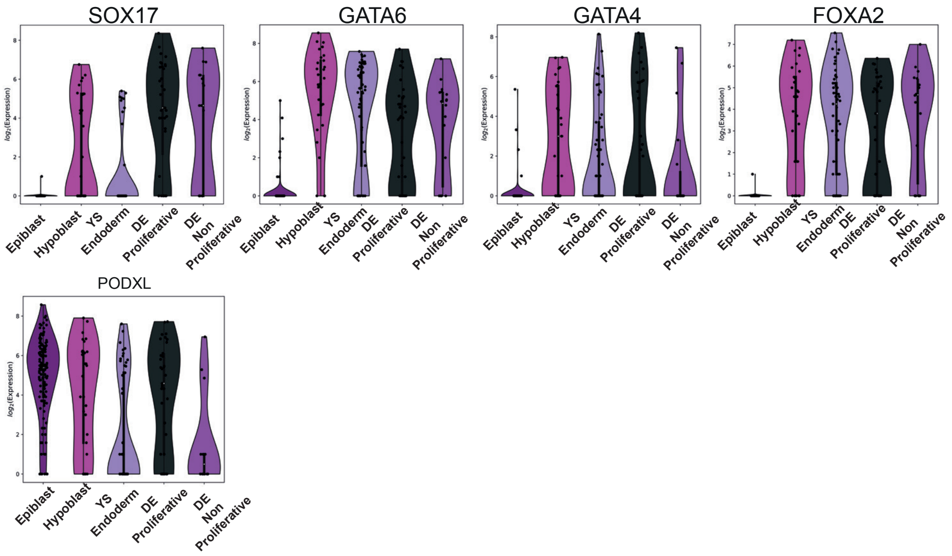


## B

##### Hypoblast markers lowly expressed in DE

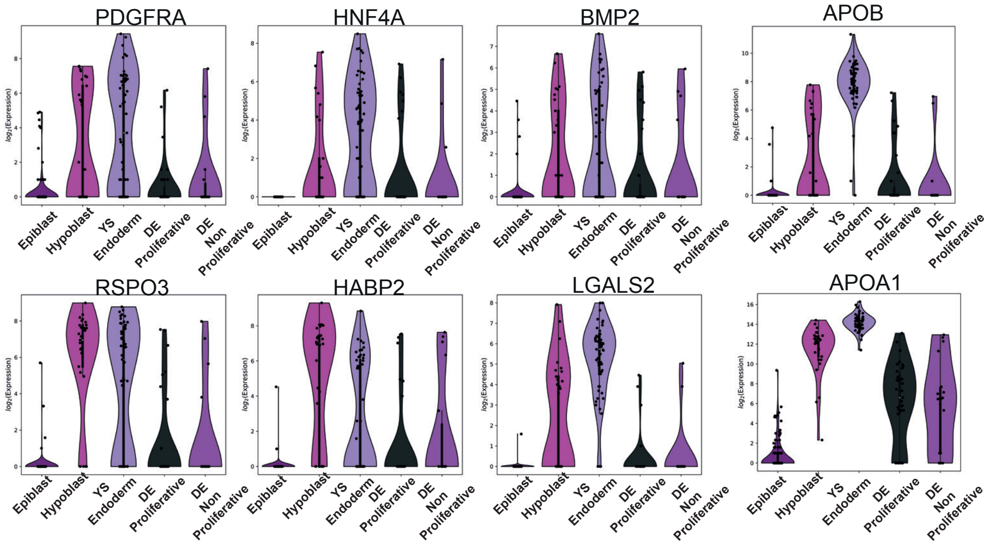


## C

##### Hypoblast markers expressed in 10X timecourse

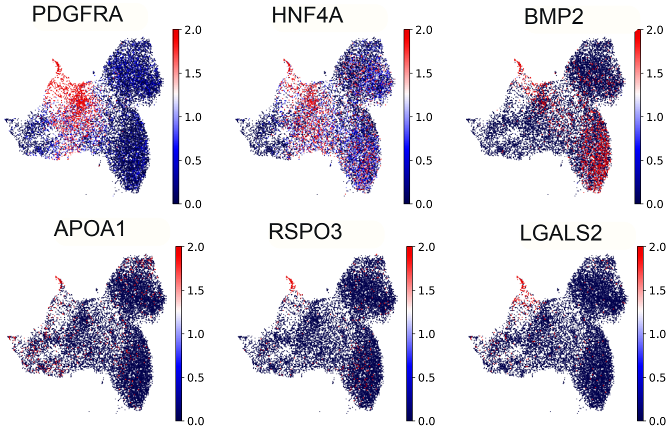

### Supplemental Figure 3

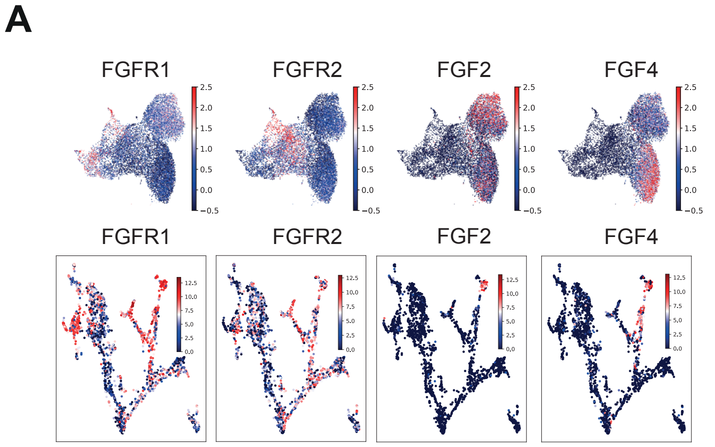


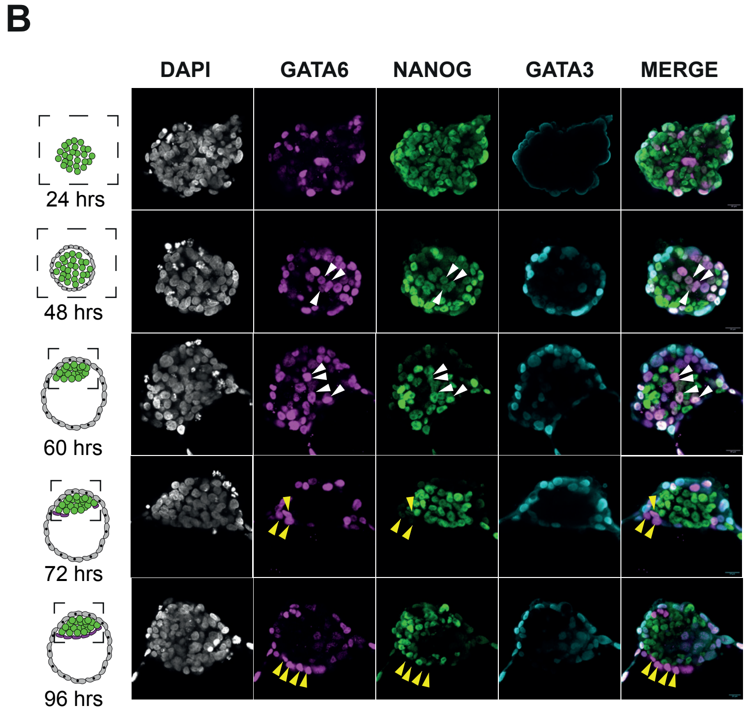

### Supplemental Figure 4

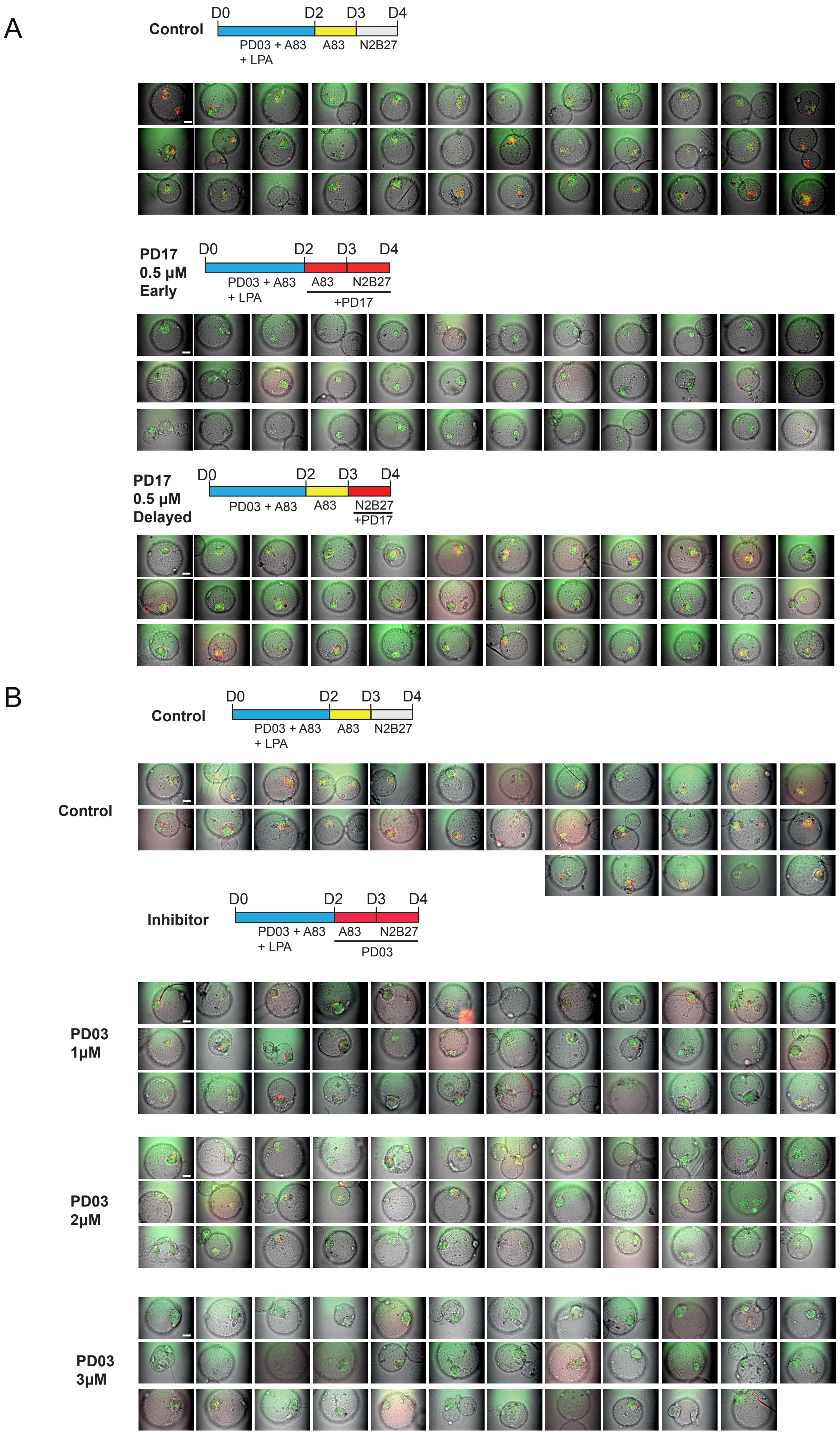

### Supplemental Figure 5

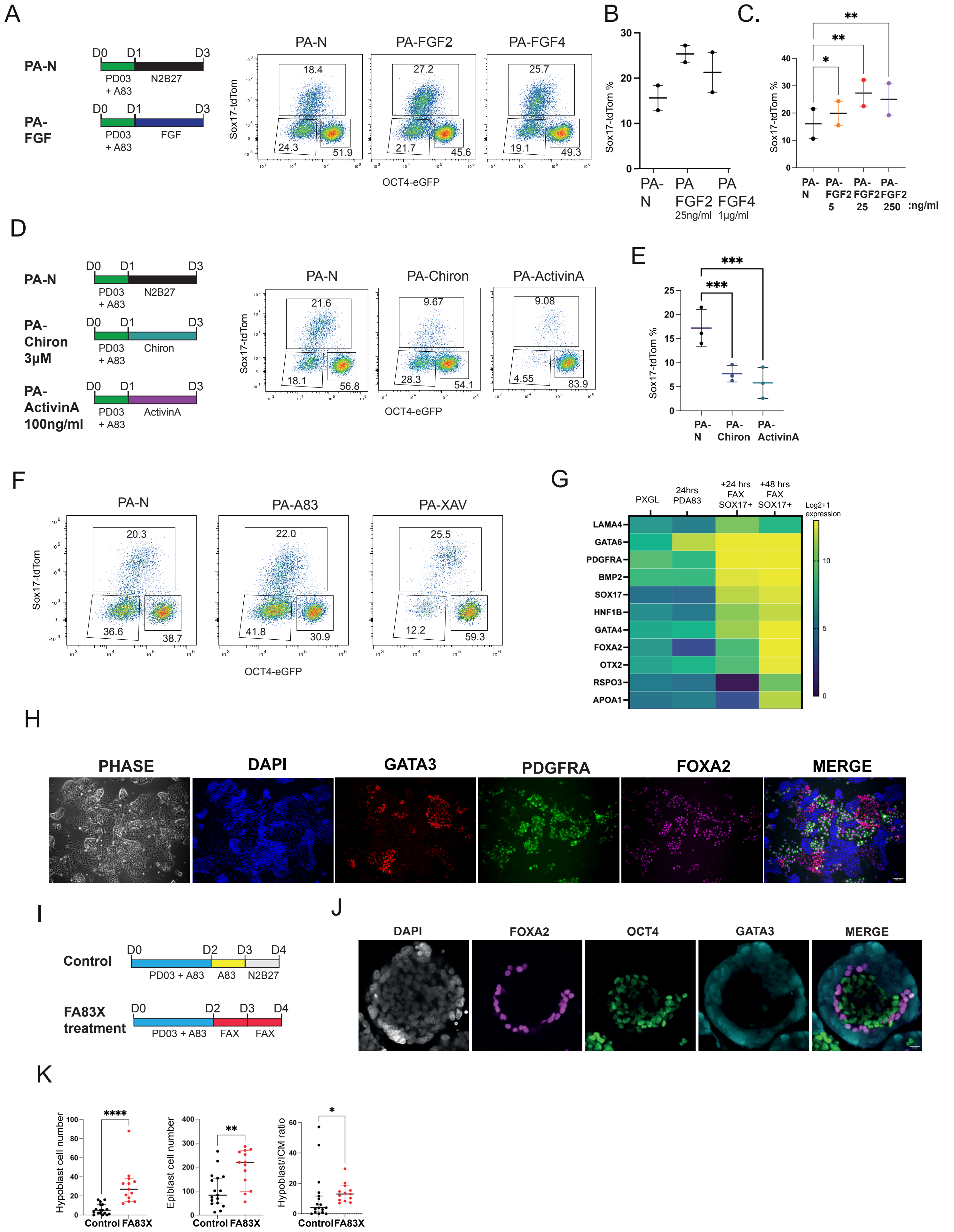

### Supplemental Figure 6

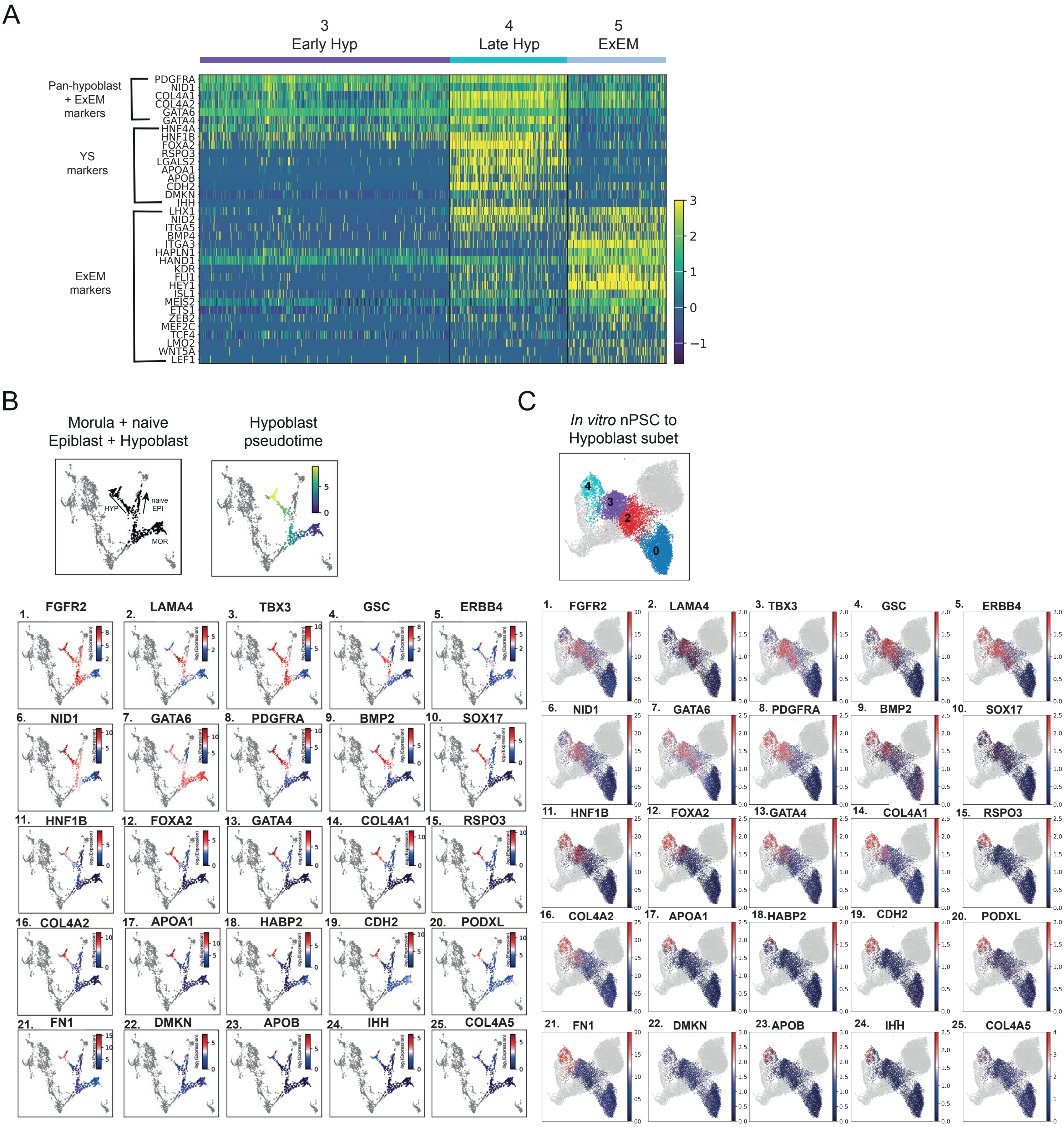
